# Pilot study of EVIDENCE: High diagnostic yield and clinical utility of whole exome sequencing using an automated interpretation system for patients with suspected genetic disorders

**DOI:** 10.1101/628438

**Authors:** Go Hun Seo, Taeho Kim, Jung-young Park, Jungsul Lee, Sehwan Kim, Dhong-gun Won, Arum Oh, Yena Lee, In Hee Choi, Jeongmin Choi, Hajeong Lee, Hee Gyung Kang, Hee Yeon Cho, Min Hyun Cho, Yoon Jeon Kim, Young Hee Yoon, Baik-Lin Eun, Robert J Desnick, Changwon Keum, Beom Hee Lee

## Abstract

**Purpose:** EVIDENCE, an automated interpretation system, has been developed to facilitate the entire process of whole exome sequencing (WES) analyses. This study investigated the diagnostic yield of EVIDENCE in patients suspected genetic disorders.

**Methods:** DNA from 330 probands (age range, 0–68 years) with suspected genetic disorders were subjected to WES. Candidate variants were identified by EVIDENCE and confirmed by testing family members and/or clinical reassessments.

**Results:** The average number of overlapping organ categories per patient was 4.5 ± 5.0. EVIDENCE reported a total 244 variants in 215 (65.1%) of the 330 probands. After clinical reassessment and/or family member testing, 196 variants were identified in 171 probands (51.8%), including 115 novel variants. These variants were confirmed as being responsible for 146 genetic disorders. One hundred-seven (54.6%) of the 196 variants were categorized as pathogenic or likely pathogenic before, and 146 (74.6%) after, clinical assessment and/or family member testing. Factors associated with a variant being confirmed as causative include rules, such as PVS1, PS1, PM1, PM5, and PP5, and similar symptom scores between that variant and a patient’s phenotype.

**Conclusion:** This new, automated variant interpretation system facilitated the diagnosis of various genetic diseases with a 51% improvement in diagnostic yield.

## Introduction

Whole exome sequencing (WES) using massively parallel sequencing techniques has identified the specific genetic defects in over 1000 Mendelian disorders during the past decade. To date, more than 7000 genetic disorders involving more than 4000 genes have been identified, and the numbers continue to increase as the genetic defects in additional disorders were identified.^1,2^ The diagnostic rate of WES has been found to range from 30% to 40%, a variation that may be attributed to the numbers and phenotypes of enrolled patients and the various characteristics of study cohorts.^3–10^

Whole genome studies such as WES are time-consuming and labor-intensive, requiring clinical geneticists and bioinformaticians to match large numbers of candidate variants with various clinical symptoms in each subject analyzed.^11^ Moreover, in the absence of supporting data, many variants remain “variants of uncertain significance” (VUS), limiting the ability to confirm genetic diagnoses.^12^

Guidelines of the American College of Medical Genetics (ACMG) attempted to prioritize genetic variants and led to the development of several bioinformatic tools.^13,14^ These tools, however, have limited ability to accurately predict the pathogenicity of each variant. Phenotype-centric interpretation methods were developed using several computational tools, which automatically prioritized the genetic variants in each patient and ranked them, according to the biological function of each gene, the molecular impact of the variant, and the relationship of the variant to that patient’s phenotype.^9,15,16^ Although these approaches noticeably reduced the number of candidate variants responsible for the disease phenotype in each patient, these numbers varied among studies, without significantly improving genetic diagnosis rates, which have remained at about 30–35%.^5,16^

This study describes a new, streamlined, automated interpretation system, termed EVIDENCE (3Billion, Inc., Seoul, South Korea), which interprets over 100,000 variants according to ACMG guidelines^17^ and prioritizes variants based on each patient’s phenotype within a few minutes. A symptom suggestion system based on Human Phenotype Ontology (HPO) was created to capture most patient phenotypes. Finally, the EVIDENCE system was able to calculate similarity scores between the clinical phenotypes suggested by the candidate variants and actual patient phenotypes, to match this score with the genetic diseases listed in the OMIM database (www.omim.org). This pilot study found that EVIDENCE significantly improved the rate of diagnosis of a variety of genetic diseases.

## Materials and Methods

### Recruitment of patients

The study enrolled 330 patients, clinically suspected of carrying a genetic disorder, from 330 non-consanguineous unrelated families, who presented at the Medical Genetics Center, Asan Medical Center, Seoul, South Korea, from April 2018 to August 2019. Their detailed demographic and clinical characteristics were reviewed, including age and diagnosis at presentation, sex, family history, laboratory findings, radiologic findings, and genetic testing results.

Patients aged ≥5 months were included if they were strongly suspected of having a genetic disease by medical geneticists and were undiagnosed, despite conventional genetic tests such as chromosome analyses, chromosome microarray, or single or targeted gene panel testing. Patients aged <5 months were included if they had a congenital anomaly in one or more major organs, including the brain, heart, or gastrointestinal, urological, or musculoskeletal systems, or if they were strongly suspected of having a genetic disease by medical geneticists or radiologists.

Informed consent was obtained from patients or their parents after genetic counseling regarding the WES test. The study was approved by the Institutional Review Board for Human Research of the Asan Medical Center (IRB numbers: 2018-0574 and 2018-0180).

### Whole exome sequencing, variant calling, and variant annotation

Blood, saliva, or buccal swab samples were collected from each patient, and genomic DNA was extracted from each sample. All exon regions of all human genes (~22,000) were captured using Agilent SureSelect kits (version C2, December 2018) and sequenced using the NovaSeq platform (Illumina, San Diego, USA). Raw genome sequencing data were analyzed with an in-house software program; these analyses included alignment to the reference sequence (original GRCh37 from NCBI, Feb. 2009) and variant calling and annotation. The mean depth of coverage was 100 X (>10 X = 99.2%).

### EVIDENCE: Prioritization of variants and symptom suggestion system

The streamlined variant interpretation software program, EVIDENCE, was developed in-house to prioritize variants based on each patient’s phenotype and to interpret these variants accurately and consistently within approximately five minutes. This system has three major steps: variant filtration, classification, and similarity scoring for patient phenotype. In the first step, allele frequency was estimated in population genome databases, including gnomAD (http://gnomad.broadinstitute.org/), 1000 Genomes (http://www.internationalgenome.org/), ESP (https://evs.gs.washington.edu/EVS/), and 3Billion, Inc. Common variants with a minor allele frequency >5% were filtered out in accordance with rule BA1 of the ACMG guidelines.^17^

In the second step, evidence of the pathogenicity of variants was obtained from disease databases, including OMIM (www.omim.org), ClinVar (https://www.ncbi.nlm.nih.gov/clinvar/), and UniProt (https://www.uniprot.org/); these factors included the function of each gene, the domain of interest, the mechanism of disease development, and its inheritance pattern and clinical relevance. The predicted functional or splicing effect of each variant and its degree of evolutionary conservation were evaluated using several *in silico* tools, including REVEL, ada, and ra score.^18,19^ The pathogenicity of each variant was evaluated according to the recommendations of the ACMG guidelines.^17^ In the third step, the clinical phenotype of each proband was transformed to its corresponding standardized HPO term and was assessed to measure the similarity with each of ~7000 rare genetic diseases.^20,21^ The similarity score between each patient’s phenotype and symptoms associated with that disease, caused by prioritized variants according to ACMG guidelines, ranged from 0 to 10. The entire process of genetic diagnosis, including processing of raw genome data, determining variant prioritization, and measuring the similarity between each phenotype and disease, was integrated and automated into a computational framework.

### Variant interpretation and confirmation

Relevant candidate variants, including VUS, based on EVIDENCE, were reviewed and selected by medical geneticists. After another examination in the outpatient clinic, the DNA of each patient and/or their parents was subjected to Sanger sequencing to confirm the initial variant.

### Statistical analysis

All statistical analyses were performed with R studio software (version 3.5.1). Principal component analysis (PCA) of symptoms and genetic variations required construction of a patient symptom matrix and a genetic variation matrix for each patient, with entries of 1 for patient j having symptom or variant i, and entries of 0 otherwise. All pathogenic variants aggregated from the entire patient cohort were used. Ten types of functional variations were treated separately, resulting in 1285 combinations of genetic and functional variants. The entries in both matrices were calculated using a custom-made program and an Eigen C++ linear algebra library, with *P* < 0.05 considered statistically significant.

## Results

### Patient demographics

The demographic characteristics of the 330 patients are shown in Table 1. Mean ages at clinical presentation and at performance of WES were 5.9 ± 12.9 years (range, 0–68 years) and 11.9 ± 16.2 years (range, 0–70 years), respectively. Of the 330 patients, 246 (74.5%) were under 18 years of age. Patients manifested a broad range of phenotypes across organ systems. The average number of systems manifesting phenotypic abnormalities per patient was 4.5 ± 5.0. Abnormalities in the nervous system were the most frequent, observed in 60% of patients, followed by the musculoskeletal system (53.9%), the head and neck (43.3%), the cardiovascular system (26.9%), and the endocrine system and metabolism (24.2%) (Table 1). Of the total 16,000 HPO terms, 550 terms were identified in these 330 patients. These terms were broadly distributed throughout the genome, with the HPO terms matching patient symptoms colored red in the Cytoscape 3.7.1 visualized network (Figure 1). These findings indicate that the phenotypes of our patients cover almost the entire range of human disease phenotypes described to date.

**Figure 1.**
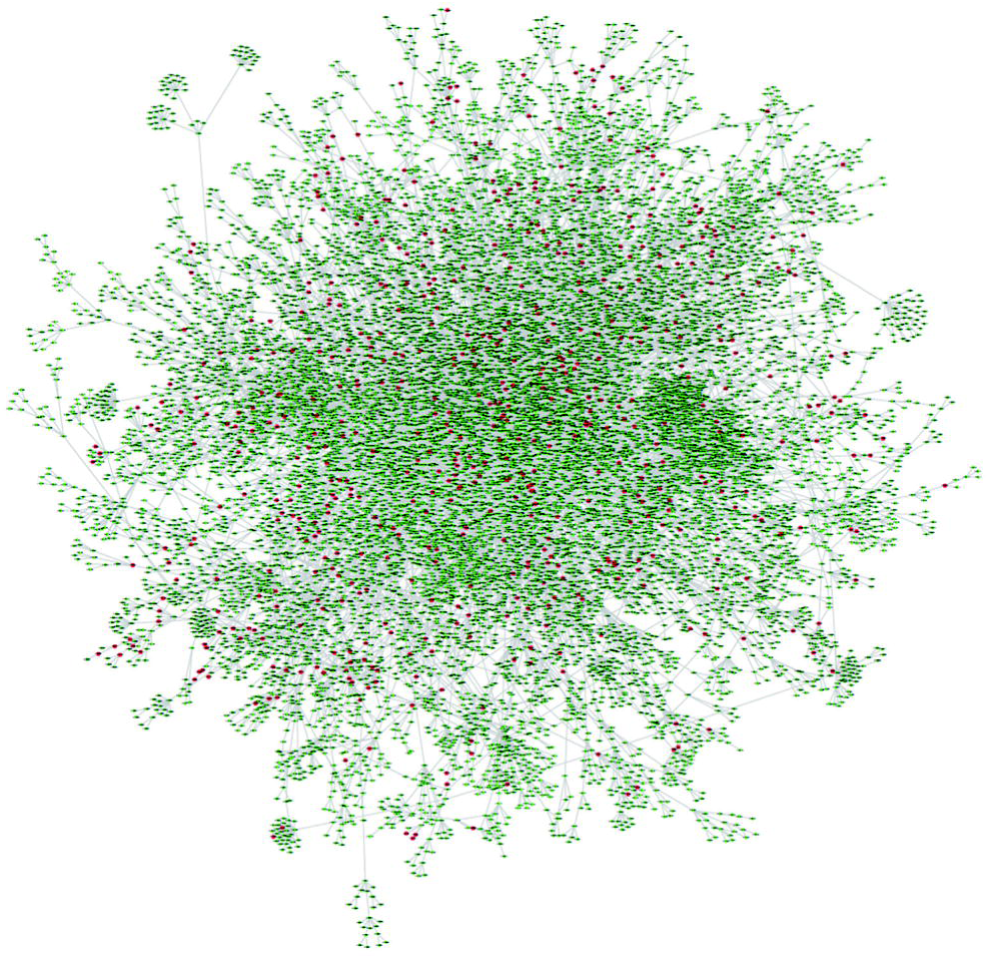
Distribution of Human phenotype ontology (HPO) terms and patient symptoms in the 330 patients (green dots: 16,000 HPO terms; red dots: patient symptoms).

Of the 330 patients, 214 (64.8%) underwent genetic testing before WES. Thirty-eight patients underwent targeted exome sequencing (including 4813 OMIM genes), and six underwent array comparative genome hybridization, respectively, with none showing diagnostic variants. Ninety-three patients underwent single gene testing for monogenic disorders. Other genetic tests included karyotyping and/or fluorescence in situ hybridization (n = 131), multiplex ligation-dependent probe amplification analyses for chromosomal microdeletion or duplication syndromes (n = 45), and mitochondrial full genome sequencing analysis (n = 20). All of the tests did not reveal a specific diagnosis in the patients tested.

### Diagnostic yield and classification of identified variants

The number of patients with variants and the identity of these variants are summarized in Figure 2. EVIDENCE identified an average of 15.0 ± 8.7 variant-disease pairs per patient, according to ACMG guidelines and similarity scores. Medical geneticists and bioinformaticians evaluated each candidate variant and selected the variant most closely associated with each patient’s phenotype.

**Figure 2.**
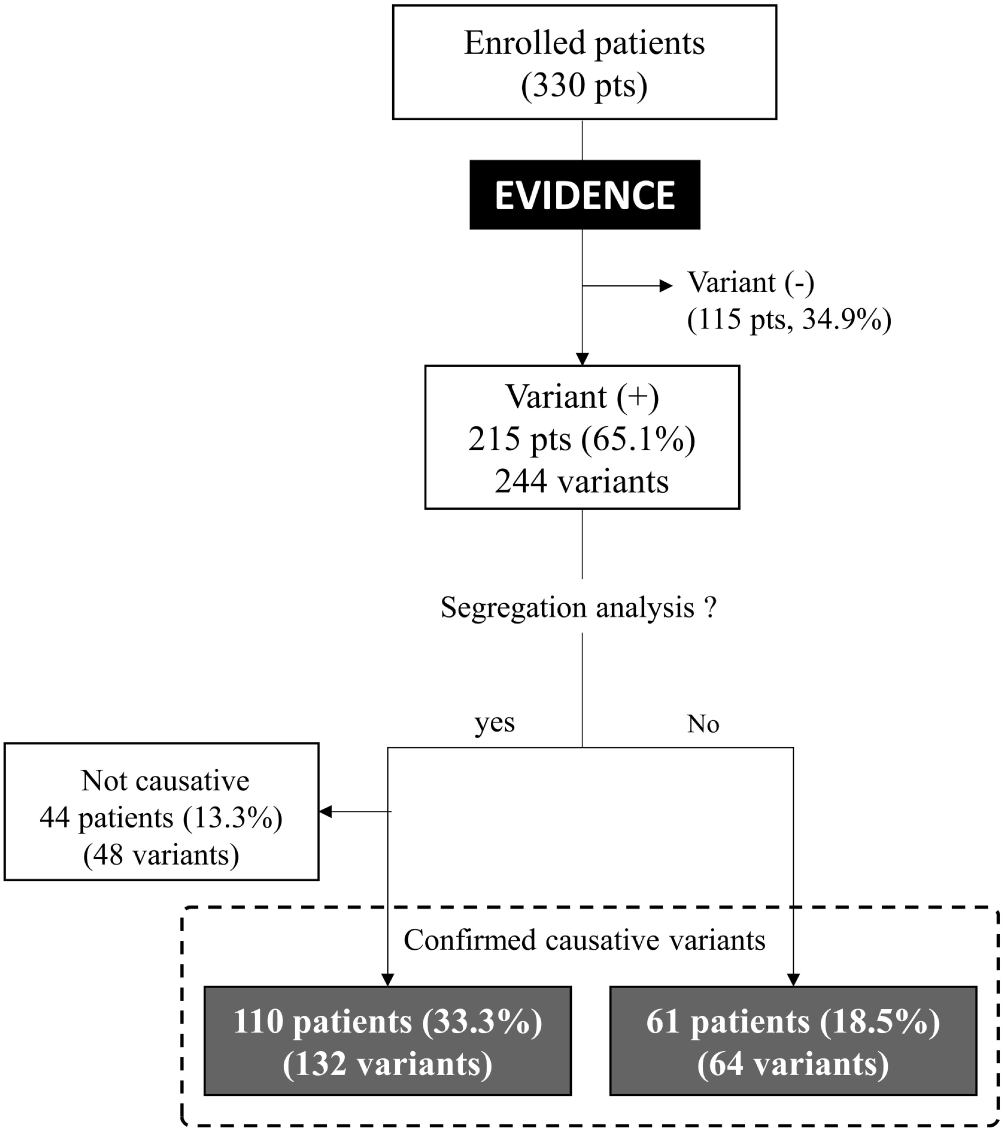
Schematic diagram showing the number of patients with and without variant identification and segregation analysis.

EVIDENCE identified 244 variants, including 131 VUS, in 215 (65.1%) of the 330 patients. Among these, 180 variants from 154 patients (46.7%) were assessed by Sanger sequencing and familial segregation analysis, with 132 variants in 110 patients (33.3%) confirmed as causative. In addition, 64 variants from 61 patients (18.5%) were confirmed based on the function of the identified gene and the predicted pathogenicity of the variant, as well its frequency in the general population. In summary, 196 variants in 171 patients (51.8%), including 115 novel variants, were confirmed as responsible for 146 genetic disorders. The remaining 48 variants in 44 patients (13.3%) were not regarded as causative because they were inherited from an asymptomatic parent, and the putative gene represented a dominant disorder with expected high penetrance or was found in cis pattern in a recessive disorder. Rates of diagnosis did not differ significantly in patients who did and did not undergo genetic testing before WES (53.3% [114/214] *vs.* 49.1% [57/116], *P* = 0.491).

The inheritance pattern of identified variants in the 171 patients was autosomal dominant (n = 120, 70.2%), autosomal recessive (n = 34, 19.9%), and X-linked (n = 17, 9.9%). Of the 196 confirmed variants, 52 (26.5%) were regarded as *de novo*, and 39 (19.9%) were assumed to be *de novo*. Forty-seven variants from 25 patients inherited in an autosomal recessive manner were detected in a trans pattern.

According to ACMG guidelines, nine (4.6%) variants were regarded as pathogenic, 98 (50%) as likely pathogenic, and 89 (45.4%) as of uncertain significance (Figure 3). After clinical assessment, including biochemical tests, imaging analysis and physical examination, 70 (35.7%) variants were regarded as pathogenic, 48 (24.5%) as likely pathogenic, and 78 (39.8%) as of uncertain significance, and then after family segregation analysis, 95 (48.5%) variants were regarded as pathogenic, 49 (25%) as likely pathogenic, and 52 (26.5%) as of uncertain significance. In total, 124 patients (37.6%) had pathogenic or likely pathogenic variants. The list of variants and diseases were described in Table S1.

**Figure 3.**
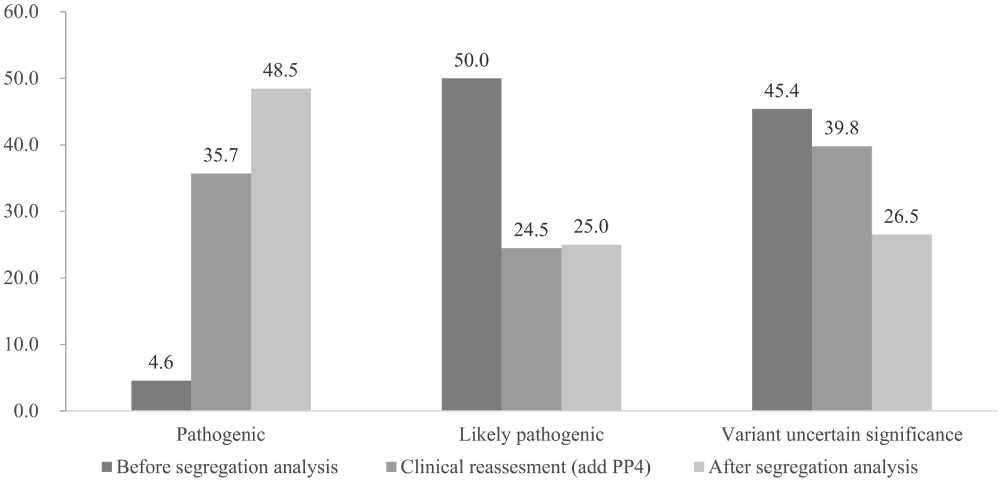
Distribution of the likely pathogenicity of identified variants by EVIDENCE before segregation analysis, after addition of PP4, and after segregation analysis.

### Characteristics of the confirmed variants

The characteristics of 196 variants confirmed to be disease-causing and the other 48 variants regarded as not being disease-causing were compared based on ACMG guidelines and symptom similarity. Of the 113 pathogenic or likely pathogenic variants, 107 (95%) were confirmed as being causative compared with 89 (67%) of the 131 VUS. Of the 48 variants regarded as not being disease-causing, 42 (87.5%) were categorized as being of uncertain significance, whereas only six (12.5%) were regarded as likely pathogenic; these six variants were found in the asymptomatic parents of a child with an autosomal dominant disorder with expected high penetrance.

Seven items in the ACMG guidelines, PS2, PS3, PS4, PM3, PM6, PP1, and PP4, can be checked after segregation analysis, functional determination, and physician assessment. Five items, PVS1, PS1, PM1, PM5, PP2, and PP5, however, had relatively high confirmation rates of >90% each (Table 2).

The average numbers of HPO items in patients without an identified variant, in those with a confirmed variant, and in those with a rejected variant were 6.4 ± 5.0, 7.4 ± 5.3, and 9.0 ± 4.9, respectively (*P* > 0.05). There was no significant difference in probability of confirmation of a certain variant identified by EVIDENCE among the affected organ types (Table S2). However, importantly, the confirmation rate was significantly higher when the similarity score of a variant was above than when it was below 5 points (*P* = 0.032, Figure 4).

**Figure 4.**
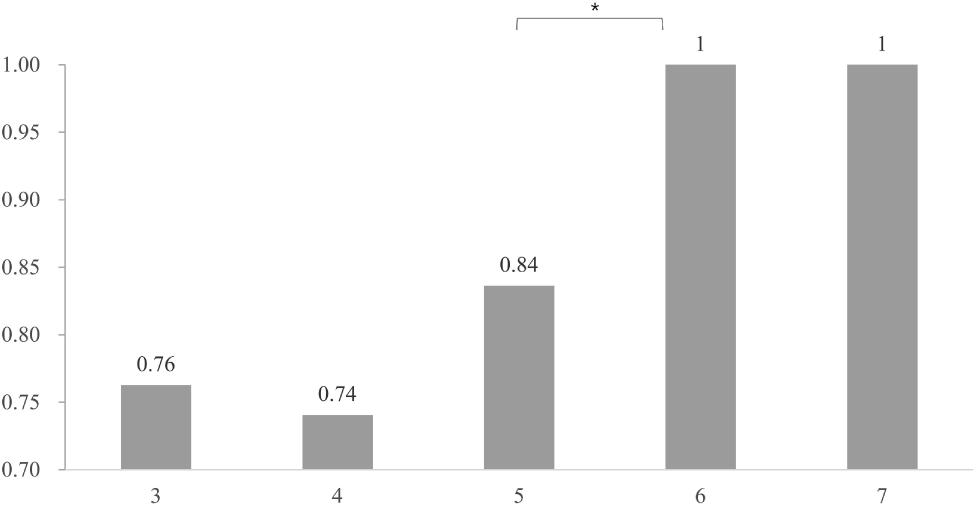
Distribution of symptom similarity scores of patient phenotypes and genetic phenotypes suggested by the automated system. * *P* < 0.05

### Genetic and phenotypic diversity of enrolled patients

No significant differences were observed in the distribution of clinical symptoms between patients with and without a variant identified by EVIDENCE (by 2-dimensional Kolmogorov–Smirov test; Figure 5A). A similarity of principal component (PC1) value in symptom PCA of two patients implies a similarity in symptoms between these patients (Figure 5B). By visual inspection of Figure 5B, we divided patients with identified variants into two groups using a PC1 of 0.5 in symptom PCA value as a threshold. Of the 215 patients with identified variants, 100 (46.5%) were clustered together in PC1 of symptom PCA ranging from 0.5 to 0.93 (13% of total symptom PCA PC1 range). That is, the phenotypes of 46.5% of the patients covered only 13% of the total symptom PCA space, with the remaining 53.5% of patients covering the other 87%. The two patient groups had similar diversities of genetic variants, as shown by a Student’s t-test of PC1 values of genetic variation PCA (*P* = 0.899).

**Figure 5.**
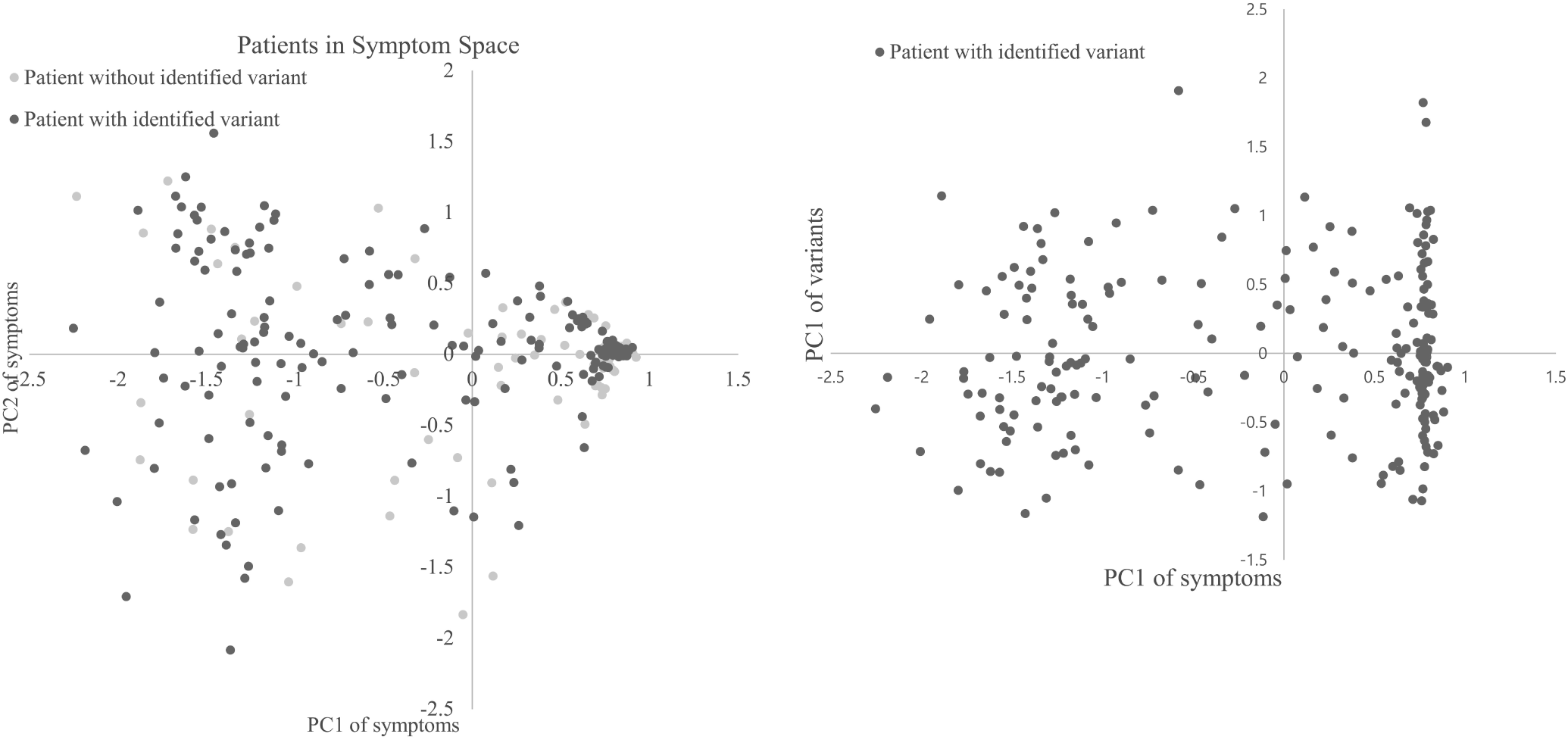
(A) Distribution of patients in symptom space. (B) Distribution of patients with identified variants in symptom and genetic variation space.

## Discussion

The EVIDENCE automated interpretation system was found to be useful in the entire WES process, including raw data processing, variant prioritization, and measurement of phenotypic similarity between patients and suggested candidate diseases. The automated workflow provided by this system reduced the total amount of time required for diagnosis from ~20–40 hours^11^ to less than 5 minutes.

The diagnostic yield of EVIDENCE in the present study (51.8%) was higher than that previously reported for automated systems (30–35%).^9,15,16^ This finding was important, as the phenotypes of the enrolled patients were quite heterogeneous, broadly dispersed, and not limited to certain organ categories. Diagnosis rates over 50% have been reported in the absence of an automated system in patients with select disease phenotypes, including hearing loss, visual impairment, or abnormalities of the musculoskeletal system, as well as in patients in critical condition and in newborns presenting with symptoms.^4,8,22^ Moreover, in the absence of an automated system, a large amount of time was required to interpret a significant numbers of variants in each patient.^6,11^ The results presented here indicate that our automated system can diagnose various types of genetic diseases with comparable accuracy, but much greater speed, than non-automated analyses.

The high rate of diagnosis achieved by the automated system may be due to its high performance efficiency. Based on the systemic analysis of each variant and the relationship of each variant to patient phenotype, the results of this analysis suggested an average of 15 variants, putatively responsible for a patient’s phenotype. This reduction in variant number shortened the time required to select the variant most likely responsible for that patient’s phenotype, and it minimized the likelihood of missing the disease-causing variant.

Another factor responsible for the high diagnostic rate of this automated system was that a substantial proportion of the variants suggested by the system were VUSs. These VUSs were subsequently tested in family member segregation analysis and phenotype reassessment, as it is unclear whether VUSs are causative variants in the absence of segregation analysis and clinical reassessment. Updated information on variants in genome databases can result in VUSs being classified as pathogenic or benign.^12,23,24^ Before family testing and clinical reassessment, 28.8% of the patients in our study had variants classified as pathogenic or likely pathogenic; after family testing and reassessment, however, 37.6% of our patients had these variants. Following segregation analysis and clinical reassessment, 37 (41.6%) of 89 VUSs were reclassified as pathogenic or likely pathogenic.

Recently, the Clinical Genome Resource (ClinGen) recategorized variants according to ACMG guidelines by focusing on unique features of particular genes or genomic regions.^25–30^ Our study applied rules of the ACMG guidelines, such as PVS1, PS1, PM1, and PM5, to variant interpretation, focusing on the characteristics of each variant, including the type of variant, gene function, and the role of gene domains. The value of applying the PP5 rule to the validation process remains unclear, but ACMG guidelines have not been updated to delete the PP5 rule.^17,31^ In our study, the PP5 rule was exclusively applied to those variants that were found in non-overlapping sources. Moreover, the elimination of this rule would not affect variant classification.

In variant prioritizing systems, the score of the top-ranked variant increases when patient symptoms more precisely match those caused by the responsible gene, and when the number of HPO terms of a patient increases.^32^ The present study found no significant differences in the average numbers of HPO terms and organ types between patients in whom causative variants have and have not been identified. This finding was probably related to the wide range of phenotypes among our patients. By contrast, similarity scores were calculated as described with little modification.^20,21^ Maximal depth of the common ancestor node of two symptoms in the HPO tree structure was used instead of its information content because the latter depends on symptom-disease mapping data. Notably, we observed that scores ≥5 points were associated with a significantly higher probability of confirmation of a certain variant as causative. Because the ancestor HPO term has relatively low accuracy, however, improvements in the determination of similarity scores and more detailed description of symptoms are required to enhance the accuracy of variant prioritizing systems.

This study had several limitations. First, most of the patients were pediatric patients. Pediatric patients have a higher likelihood of genetic diseases than adults, which may have contributed to the relatively high rate of diagnosis in our patient cohort. Second, segregation analysis could not be performed in 60 of the 171 patients diagnosed with genetic diseases because samples from family members were unavailable. Family testing of all 60 of these patients may have altered the diagnosis rate between 33% and 54%. Third, the pathogenicity of VUS can be altered by updates in variant information. Finally, the actual causative variant may have been missed by the automated system.

In conclusion, the rate of detection of variants by the automated system did not differ significantly in patients who did and did not undergo genetic testing before WES. This automated system achieved a high diagnostic yield in patients with a broad range of genetic diseases, suggesting that WES may be one of the first diagnostic methods used in patients suspected of having a genetic disease, and that the automated system can facilitate the diagnostic process. This new method will be available to others (December, 2019, https://3billion.io/) and more advanced and updated analytic tools will allow the efficiency of this system to be evaluated in a larger patient cohort.

## Acknowledgments

This work was supported by an Institute for Information and Communications Technology Promotion (IITP) grant funded by the Korean government (MSIT) (2018-0-00861, Intelligent SW Technology Development for Medical Data Analysis).

## Authors’ Contributions

BHL designed the study. GHS, TK, RJD and BHL drafted the manuscript and analyzed the data. GHS, AO, YL, IHC, JC, HL, HGK, HYC, MHC, YJK, YHY, BE, and BHL treated the patients and performed all clinical analyses. GHS, SK, JL, DW, and CK developed EVIDENCE. GHS, JYP, and TK performed the genetic analyses and interpreted variants. All authors were involved in analyzing and interpreting data. All authors read and approved the final manuscript.

---

Supplementary Table 1. Detailed information about the 196 confirmed variants in 171 patients

Supplementary Table 2. Confirmation rate according to the type of involved organ of patients with confirmed and rejected variant

## Notes

#### Summary of Updates

We revised contents about results and authors

