## Supplementary material for "Pilot study of EVIDENCE: High diagnostic yield and clinical utility of whole exome sequencing using an automated interpretation system for patients with suspected genetic disorders": Table 1

Table 1. Demographic and clinical characteristics of the patient cohort

| Category | Number (%) |
| --- | --- |
| Sex (male:female) | 164:166 (49.7:50.3%) |
| Age at presentation, years | 5.9 ± 12.9 (range, 0–68) |
| Age at time of whole exome sequencing, years | 11.9 ± 16.2 (range, 0–70) |
| Average number of overlapping phenotypes | 4.3 ± 5.0 |
| Nervous system, including behavior and/or cognition  Head or neck, including facial dysmorphism  Eye system  Ear system  Cardiovascular system  Respiratory system  Gastrointestinal system  Genitourinary system  Endocrine and metabolism/homeostasis system  Musculoskeletal and limb system  Connective tissue system  Blood and immune system  Skin  Neoplasm  Growth  Abnormality of prenatal development or birth | 198 (60%)  143 (43.3%)  79 (23.9%)  62 (18.8%)  89 (26.9%)  17 (5.2%)  56 (16.9%)  76 (23%)  80 (24.2%)  178 (53.9%)  21 (6.4%)  57 (17.3%)  56 (16.9%)  29 (8.8%)  74 (22.4%)  42 (12.7%) |
| Previous genetic analysis | 214 (64.8%) |
| Karyotype  Fluorescence in situ hybridization  Multiplex ligand dependent probe amplification  Array comparative genome hybridization  Single gene test  Targeted exome sequencing or panel test  Mitochondrial full genome sequencing | 123 (37.3 %)  8 (2.4%)  45 (13.6%)  6 (1.8%)  93 (28.2%)  38 (11.5%)  20 (6.1%) |

Results presented as mean ± SD or as number (%).
