## Supplementary material for "Pilot study of EVIDENCE: High diagnostic yield and clinical utility of whole exome sequencing using an automated interpretation system for patients with suspected genetic disorders": Table 2

Table 2. Confirmation rate according to American College of Medical Genetics guideline rules of patients with confirmed and rejected variants

| Rules | Total variants | Confirmed variants | Rejected variants | Confirmation rate |
| --- | --- | --- | --- | --- |
| PVS1 | 57 | 54 | 3 | 0.947 |
| PS1 | 44 | 42 | 2 | 0.955 |
| PM1 | 63 | 58 | 5 | 0.921 |
| PM2 | 214 | 174 | 40 | 0.812 |
| PM4 | 7 | 3 | 4 | 0.429 |
| PM5 | 13 | 12 | 1 | 0.923 |
| PP2 | 1 | 1 | 0 | 1 |
| PP3 | 110 | 93 | 17 | 0.845 |
| PP5 | 18 | 18 | 0 | 1 |
| BP1 | 3 | 2 | 1 | 0.667 |
| BP4 | 59 | 38 | 21 | 0.644 |
